# Testing and Validation of Reciprocating Positive Displacement Pump for Benchtop Pulsating Flow Model of Cerebrospinal Fluid Production and Other Physiologic Systems

**DOI:** 10.1101/2021.12.26.474197

**Authors:** Ahmad Faryami, Adam Menkara, Daniel Viar, Carolyn A Harris

## Abstract

**Background:** The flow of physiologic fluids through organs and organs systems is an integral component of their function. The complex fluid dynamics in many organ systems are still not completely understood, and in-vivo measurements of flow rates and pressure provide a testament to the complexity of each flow system. Variability in in-vivo measurements and the lack of control over flow characteristics leave a lot to be desired for testing and evaluation of current modes of treatments as well as future innovations. In-vitro models are particularly ideal for studying neurological conditions such as hydrocephalus due to their complex pathophysiology and interactions with therapeutic measures. The following aims to present the reciprocating positive displacement pump, capable of inducing pulsating flow of a defined volume at a controlled beat rate and amplitude. While the other fluidic applications of the pump are currently under investigation, this study was focused on simulating the pulsating cerebrospinal fluid production across profiles with varying parameters.

**Methods:** Pumps were manufactured using 3D printed and injection molded parts. The pumps were powered by an Arduino-based board and proprietary software that controls the linear motion of the pumps to achieve the specified output rate at the desired pulsation rate and amplitude. A range of 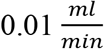 to 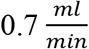 was tested to evaluate the versatility of the pumps. The accuracy and precision of the pumps’ output were evaluated by obtaining a total of 150 one-minute weight measurements of degassed deionized water per output rate across 15 pump channels. In addition, nine experiments were performed to evaluate the pumps’ control over pulsation rate and amplitude.

**Results:** volumetric analysis of a total of 1200 readings determined that the pumps achieved the target output volume rate with a mean absolute error of -0.001034283 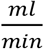 across the specified domain. It was also determined that the pumps can maintain pulsatile flow at a user-specified beat rate and amplitude.

**Conclusion:** The validation of this reciprocating positive displacement pump system allows for the future validation of novel designs to components used to treat hydrocephalus and other physiologic models involving pulsatile flow. Based on the promising results of these experiments at simulating pulsatile CSF flow, a benchtop model of human CSF production and distribution could be achieved through the incorporation of a chamber system and a compliance component.

## Background

Hydrocephalus describes a variety of conditions often presented with similar clinical symptoms. The classic presentation of hydrocephalus involves elevated intracranial pressure (ICP) which manifests with clinical symptoms such as severe headaches, visual problems, vomiting, loss of consciousness, and in extreme cases if left untreated, coma and death. Although the first experimental studies on hydrocephalus were conducted more than a century ago, the underlying mechanisms are still an active area of research [1–4].

Cerebrospinal fluid (CSF) is a blood filtrate and is 99% composed of water. CSF also contains important serum proteins such as fibrin, fibrinogen, IgG, and albumin [5]. Other than sharing biochemistry with the cerebral vasculature, flow-sensitive magnetic resonance imaging has demonstrated that CSF flow within the ventricles follows similar pulsatile flow dynamics. It is also hypothesized that an increase in intraventricular pulsations occurs as the result of changes in intracranial compliance. Many factors have been suggested to have an impact on intracranial compliance, which also determines ICP dynamics in the patient cranium [6–8].

While overproduction of CSF may result in an increase in ICP and hydrocephalus manifestation, reduction or obstruction in CSF removal and reabsorption hydrocephalus is another common pathway for the pathologic manifestation[2,4]. Another imperative difference between hydrocephalus patients is the length of time for hydrocephalus to develop and appear clinically. The clinical appearance of hydrocephalus is easily apparent in neonatal patients which involve a precipitous increase in head circumference while the pathology of hydrocephalus might take significantly longer to appear in adult patients. In some instances, the gradual progression of the disease allows that the intracranial compliance to adapt to the underlying causes of hydrocephalus rendering no clinical presentations, symptoms, or high ICP for a long time which may present clinically later in the patients’ life [9–11]. Recent studies indicate that idiopathic normal pressure hydrocephalus might be more common than it was previously supposed: new evidence suggests it appears in 5.9% of individuals older than 80 years. Studies also suggest that normal pressure hydrocephalus is underdiagnosed and undertreated due to its pathophysiology that might expand over a few decades with similarities to diseases such as Alzheimer’s and dementia. It is estimated that 3.4 per 100,000 adults undergo a surgical procedure for hydrocephalus with normal pressure hydrocephalus, high-pressure hydrocephalus, and aqueduct stenosis accounting for 47, 27, and 15 percent of the patients respectively [12].

Shunt insertion is currently the most prevalent mode of treatment for the hydrocephalic patient[13]. Nearly all patients, 98 percent, that received shunt placement therapy will experience shunt failure which involves subsequent revision surgeries, incurring up to 1$ billion in cost for hydrocephalus care [14]. Therefore, hydrocephalus diagnosis and estimation of shunt success rate for patients are among the most prolific fields of study. Flow sensitive magnetic resonance imaging is one of the diagnostic tools that has been utilized to diagnose and evaluate shunt success rates in patients. However, some of these methods are less accurate across various patient cohorts. In situ, Magnetic Resonance (MR) measurements are resource and time-intensive [15–17]. Furthermore, certain shunt systems have limited MR compatibility that could further restrict access to post-surgery MR measurements.

Alternatively, animal studies are often used to investigate CSF dynamics and to provide insights into the healthy and pathological conditions of living organisms. However, limitations of in-vivo measurements, dramatic differences between humans and animals, and the lack of control over flow dynamics leave a clear need for testing and evaluation of current modes of treatments [18]. Furthermore, simulating CSF production rate, pulsations and amplitudes might provide valuable insights into the healthy and diseased states with significant diagnostic potential.

Due to its unique anatomy and function, up to 15 percent of the cardiac output is supplied to the brain during the systole cycle and cranial blood flow and CSF are closely associated [19]. Particle tracing also demonstrated that the fluid dynamics and interactions between the microvasculature and CSF spaces are complex and region-specific [20]. One of the advantages of this pump is the capability to induce microfluidic flow rates at the physiologic beat rate and amplitude directly, negating the need to modify the system or fluid pathway to achieve a variety of flow patterns.

The main challenge of studying the fluid dynamics of hydrocephalus in a benchtop model is the complex and diverse pathology of the underlying causes that present similar clinical symptoms. Therefore, the pump must be capable of inducing pulsatile flow over a range of flow rates with accurate control over beat rate and amplitude.

## Materials and Methods

### Reciprocating Positive Displacement Pump

Pumps are comprised of five syringes, 3D printed parts, a stepper motor, a stepper motor coupler, a lead screw, linear bearings, and a series of check valves. Figure 2 illustrates a fully assembled pump. Flow into and out of the syringes is through silicone tubes. Luer tapper fittings were used to establish fluid connections between all the components. The 3D printed parts were manufactured using Anycubic I3 Mega printer using polylactic acid plastic (Anycubic Technology CO., Limited, HongKong). The stepper motor, the motor coupler, and the lead screw (composing a “drive unit”) are controlled by an Arduino-based board connected to a computer with proprietary custom-built software built using Python that determines the rotation duration and speed depending on the user-specified profile.

A user interface was developed to improve the user experience such that the pump output bulk volume rate, beat rate, and amplitude can be easily input and modified as a function of time. The rotational motion of the stepper motor is converted into linear actuation via the lead screw and linear rods and bearings. A series of check-valves are placed immediately after the syringes to allow exact volume output on the forward stroke and automatic refill on the backward motion. Each board has a capacity for up to three separate pumps for a total of fifteen channels per device.

Previously, peristaltic pumps were used to induce pulsating flow for bioreactor applications. Operators such as Flow Limiting Operator were able to successfully control the peristaltic pump output by controlling the motor rotations per minute. Inherently, peristaltic pump output and pulse rate are locked together with dependence on tubing diameter [21]. One of the main enhancements of this pump over previous designs is its capability to maintain a constant volume rate at various beat rates and amplitude.

### Data Acquisition

Flow patterns were obtained using six Sensirion SLF04 and SLF06 series flow sensors (Sensirion AG, Switzerland). The flow data were acquired using Sensirion USB Sensor Viewer software on a Hewlett-Packard desktop computer running Windows 10Pro operating system. The flow sensors were placed immediately after the output check valve to measure the pump output directly. The amplitude and beat rates were measured using the flow sensors across six separate channels, two channels per pump. While each pump is comprised of five syringes that are mounted on the same linear actuator, the flow through each channel was individually recorded and analyzed to ensure a consistent flow pattern and output through the channels on the same pump and across all three pumps connected to the same device.

### Volumetric Analysis

Although the physiologic rate of CSF production and absorption is still an active area of research, 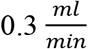 is currently accepted as the average or near average CSF production rate in a healthy adult human. A range of 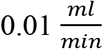 to 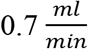 was tested to evaluate the versatility of the pumps. An arbitrary systole time of 0.3 seconds and a pulsation rate of 100 beats per minute were chosen for these experiments. The accuracy and precision of the pumps’ output were evaluated by obtaining a total of 150, one-minute weight measurements of degassed, deionized water across 15 pump channels per volume profile. The density of deionized water was measured and verified before the experiment at 1 gram per 1 milliliter of water at room temperature using a graduated beaker and an analytical balance. A Mettler Toledo AT261 DeltaRange Analytical Balance (Mettler-Toledo, LLC, USA) and Ae Adam Analytical Balance Model AAA 160DL (Adam Equipment Inc, USA) were used for all weight measurements. The flow of information between various components of the system relative to the unidirectional fluidic flow from source to output collection beakers is summarized in Figure 1. To evaluate the volumetric consistency of the pump output, the pumps were set using the user interface to run each volume rate for 10 consecutive minutes. The output of each channel was collected and weighed in individual beakers. The pumps were properly cleaned and primed using the built-in feature within the user interface before the experiments.

**Figure 1.**
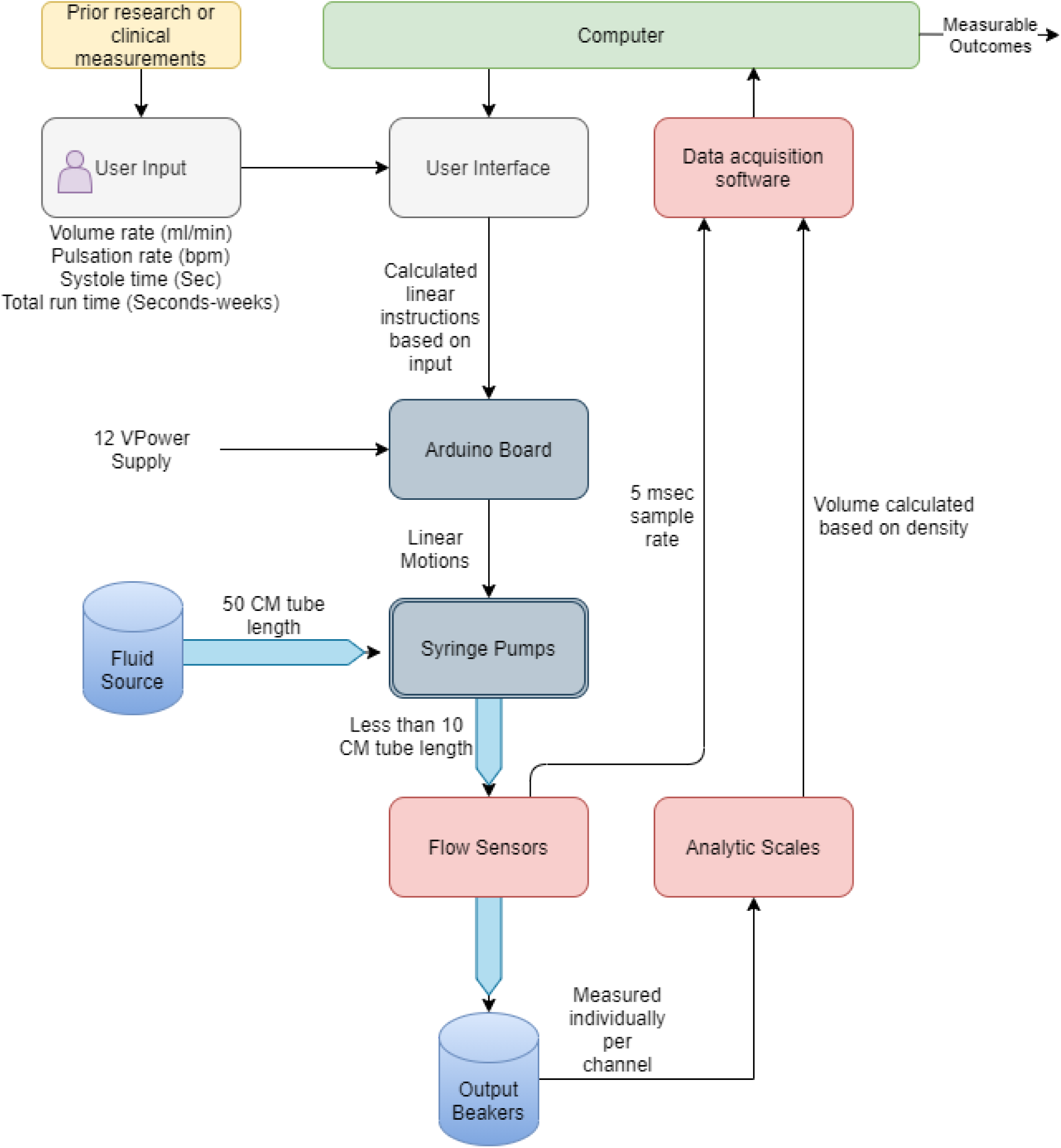
Schematic diagram illustrating the flow of information and fluids from user input to analyzed measurable outcomes

### Amplitude

Due to its implications in diagnostic and clinical applications, peak amplitude remains an important variable that was incorporated into the benchtop model. While the pump net output was solely a function of the distance the plungers were pushed forward, the amplitude of each beat was a multifactorial variable that was a function of pumps’ volume rate, beat rate, and the time required for a beat to be completed, analogous to systole time during a regular heart cycle.

The amplitude of individual beats was a function of the three variables independently specified within the user interface. According to Equation 1, the amplitude is inversely correlated to systole time and beat rate, while it is directly correlated to pump output volume rate. The dampening factor is a function of the overall compliance and material properties of all the components within the fluidic circuit including syringes, silicone tubes, and valves.

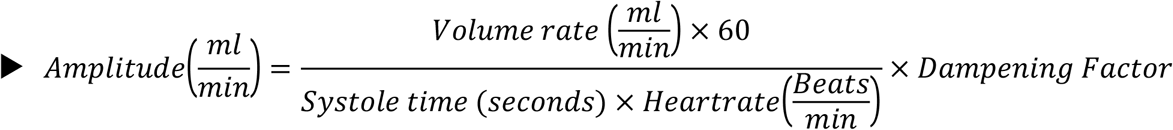

### Systole Time

Systole time is also an important variable within the domains of diagnostic and clinical applications. Systole time in the benchtop model was representative of the total duration of motor output, or the length of time the motor was in motion analogous to ventricular systole during a regular cardiac cycle. The incorporation of this variable into the benchtop model allowed for accurate control throughout the beat cycle length and inherently, the amplitude characteristics of a certain inputted profile. The manipulation of this variable across different flow profiles did not change the output volume of the profile, because this variable is an operator in the time domain and not in the volumetric output domain. Beat cycle duration was the total length of time for a complete beat cycle which was ideally equal to twice the specified systole time in a setup with no compliance. However, the compliance of silicone tubing, check valves, and residual air bubbles may have an impact on the beat cycle duration.

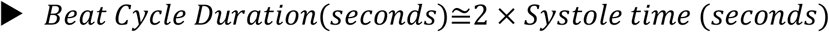

### Tubing and Valve System

One of the main considerations in the early stages of the development of the pump was its compatibility with in-vitro bioreactor projects that often require more separate channels to eliminate cross-contamination between individual samples. Therefore, each channel has a separate input and output 50 cm 3mm inner diameter silicone tube. Two check valves are implemented per channel to allow the syringes to prime and refill automatically. The mechanical inefficiencies of the system were also accounted for within the controlling program. The check valves are also useful in the automatic sterilization process of the pump and tubing system using built-in 20-minute rinse cycles with 99% isopropyl alcohol followed by three deionized water cycles to remove the remaining isopropyl alcohol from the system. Figure 2 shows the pumps from various angles with the CAD models appearing on the left and the assembled pump on the right

**Figure 2.**
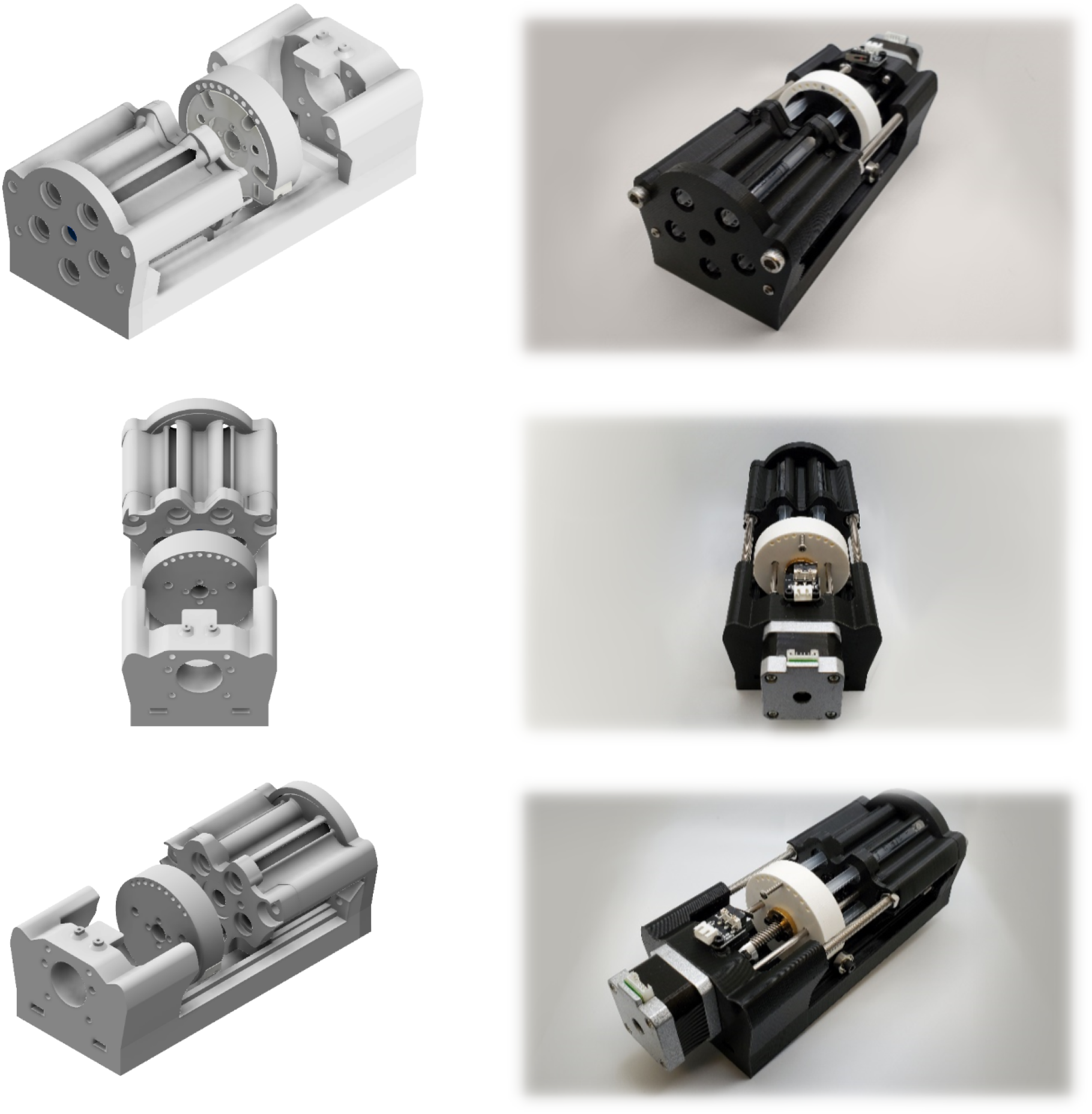
Syringe pump unit CAD models on the left. A fully assembled unit with 3D printed parts and other components

## Results

The pumps’ minimum and maximum output volume rate, beats, and amplitude limits were determined through trial and error. The optimal domain of operation for the pumps is summarized in Table 1. The accepted physiologic range for each category is included to illustrate that the non-pathologic human CSF production falls within the pumps’ optimal operation range. It is noteworthy to mention that the mechanical and software limits of the setup were not reached during these experiments. However, the reported values in Table 1 correspond to the domain at which the pumps were at peak performance.

**Table 1.**
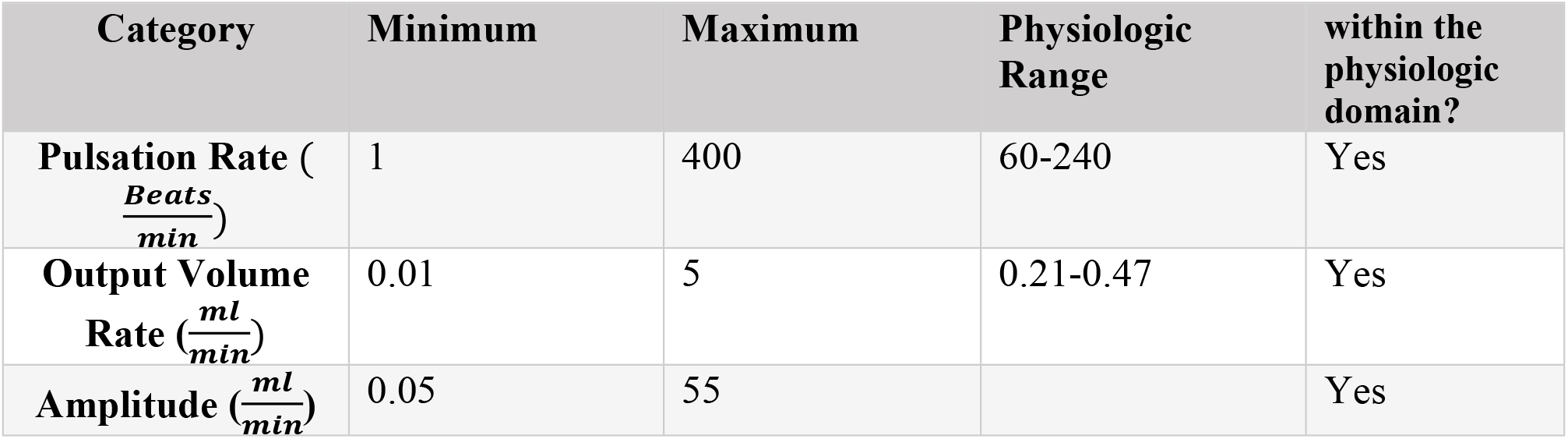
Pump optimal domain and relative to the human physiologic range

### Volumetric Analysis

The mean of 10 1-minute measurements across all fifteen channels for each specified volume rate is reported in Table 2. The highest and the lowest recorded values are also specified as well as the mean absolute error (MAE), in 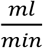 and the mean relative error in percentage. The measured mean of each output volume rate relative to the specified input is visually represented in Figure 3. The linear regression function with an *R*^2^ value of 0.9998 also indicates the accuracy and precision of the pumps over the optimal domain of operation, 0.01 to 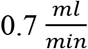 range.

**Table 2.**
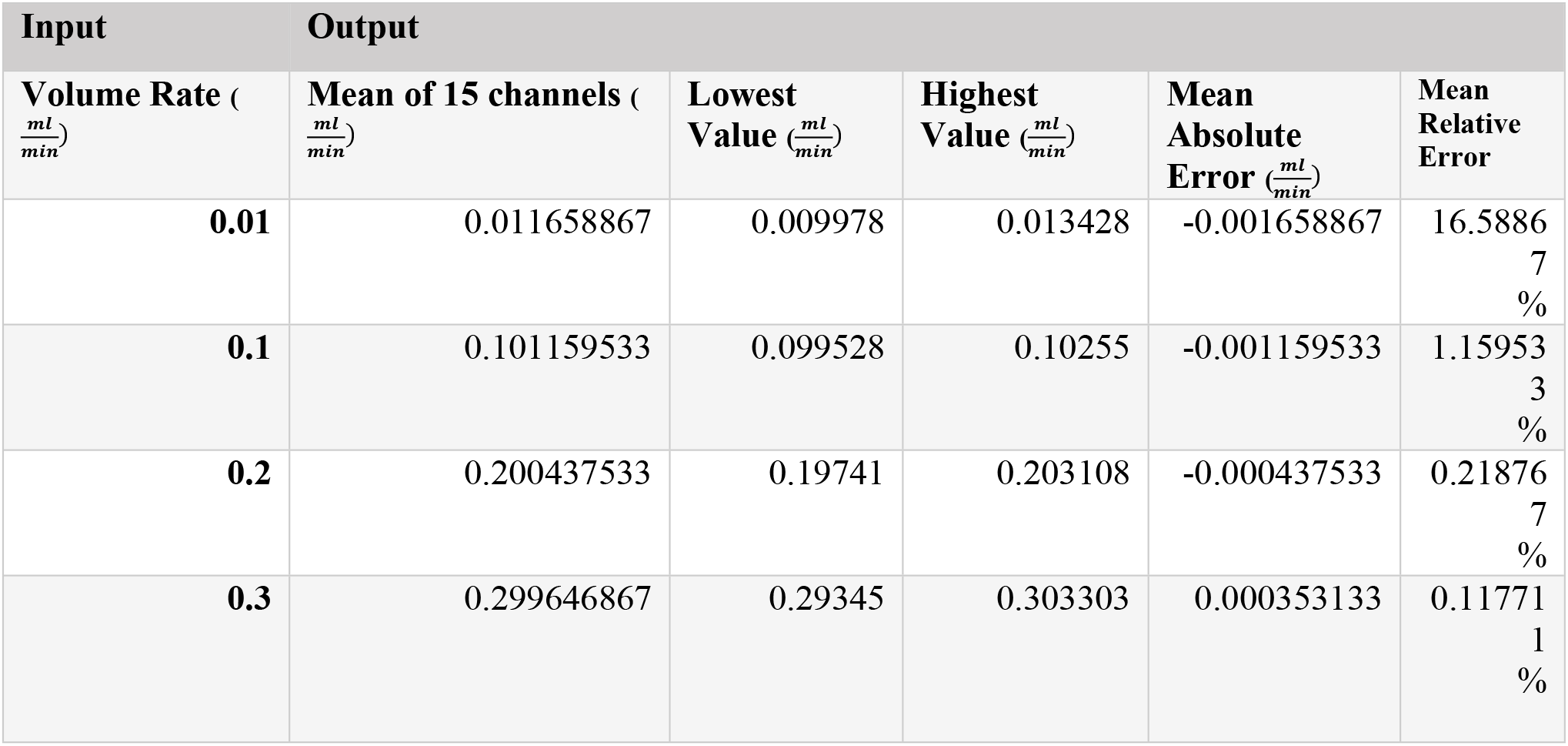
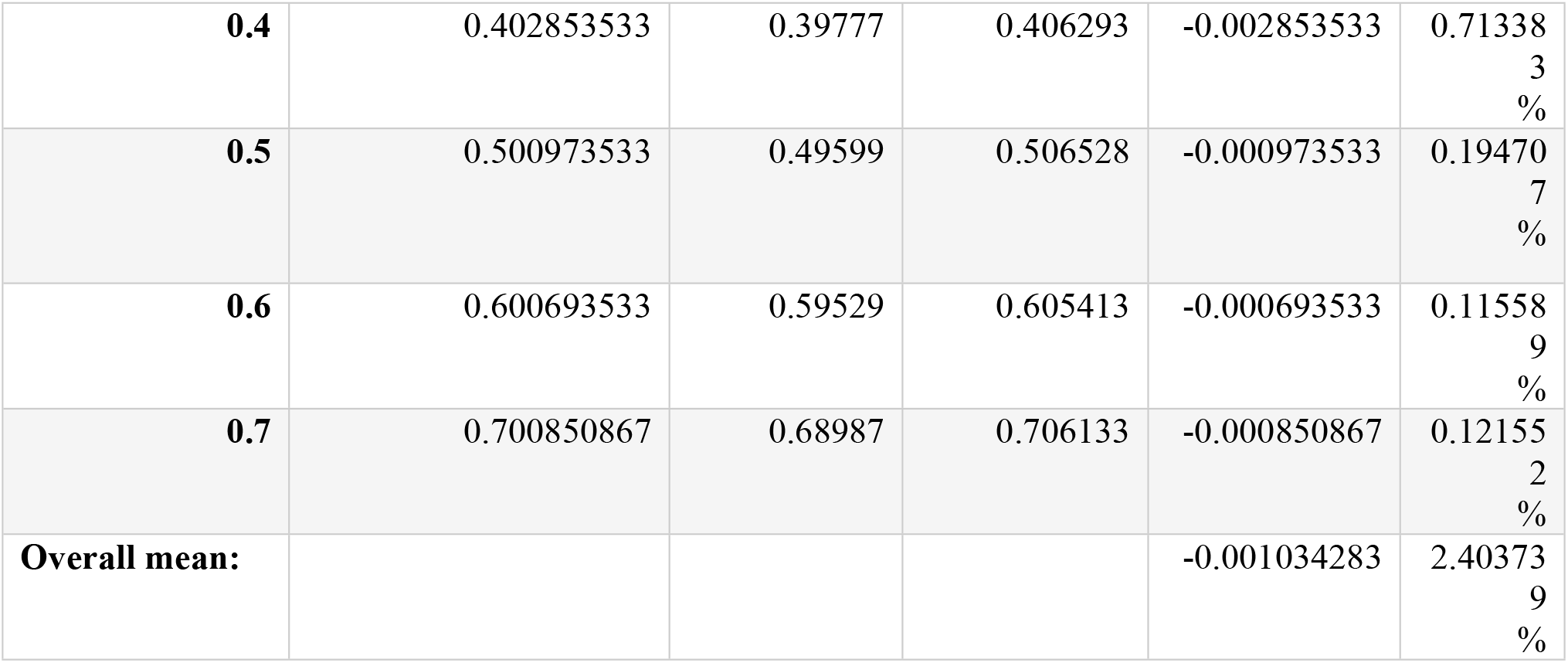
Volumetric analysis results based on weight measured by analytic balances across the 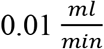 to 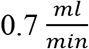

**Figure 3.**
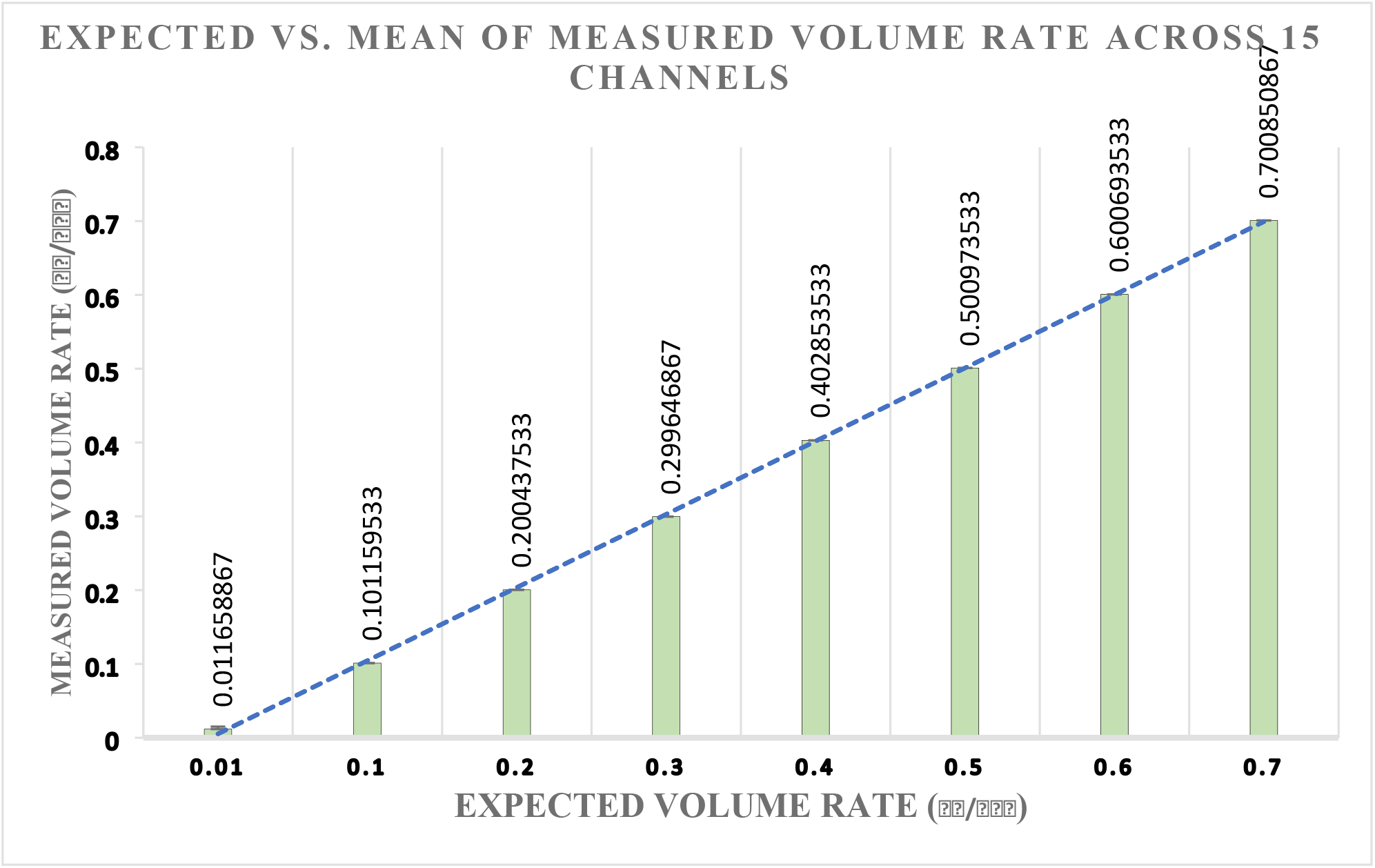
Illustrates the expected versus mean of measured volume rate across 15 channels with a linear function showing a regression

### Independent Variables

A series of tests were performed to validate the pulsatile pump output including the nine experiments depicted in Figures 4,5,6. The flow patterns were collected across 6 channels using Sensirion liquid flow sensors. On the left, the figures illustrate the pump output in 20-second segments while running at different user-specified variables, included below each set of figures. A zoomed-in 1-second interval to better illustrates the unique pattern of individual beats.

**Figure 4.**
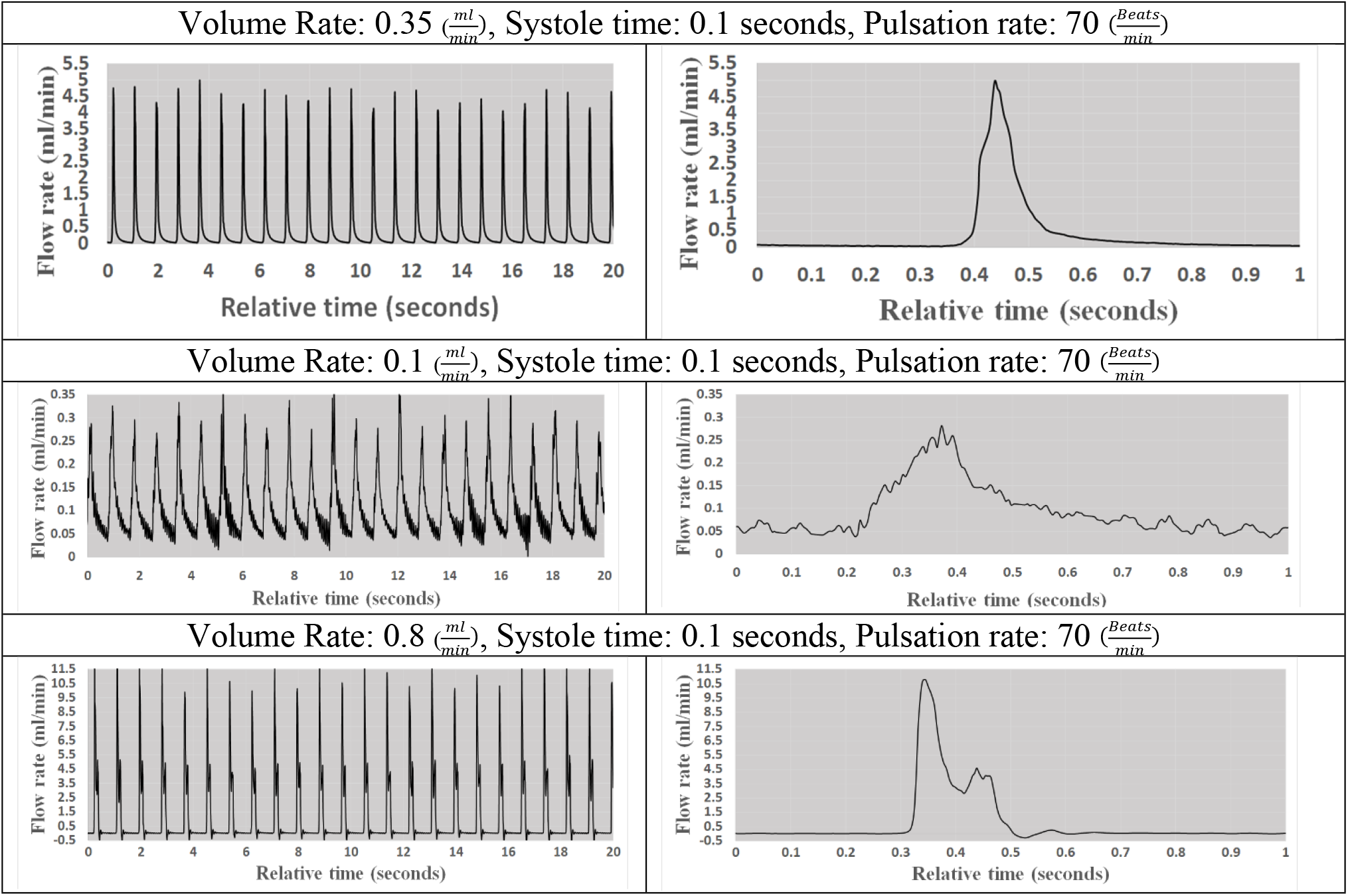
Illustrates the flow pattern at different flow rates while systole time and pulsation rate are constant

**Figure 5.**
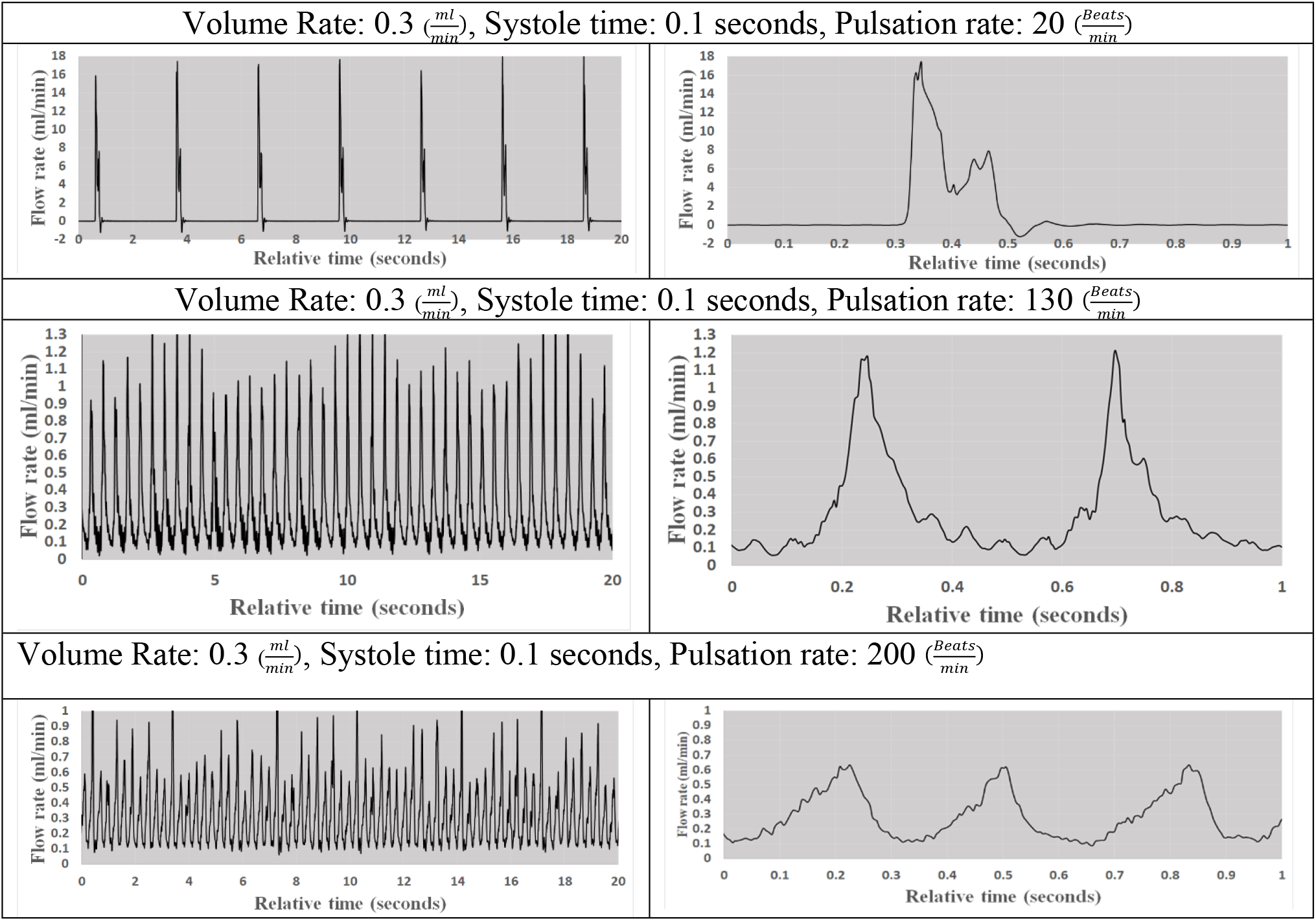
Visual representation of flow pattern at 78,200, and 20 beats per minute while volume rate and systole time remained constant

**Figure 6.**
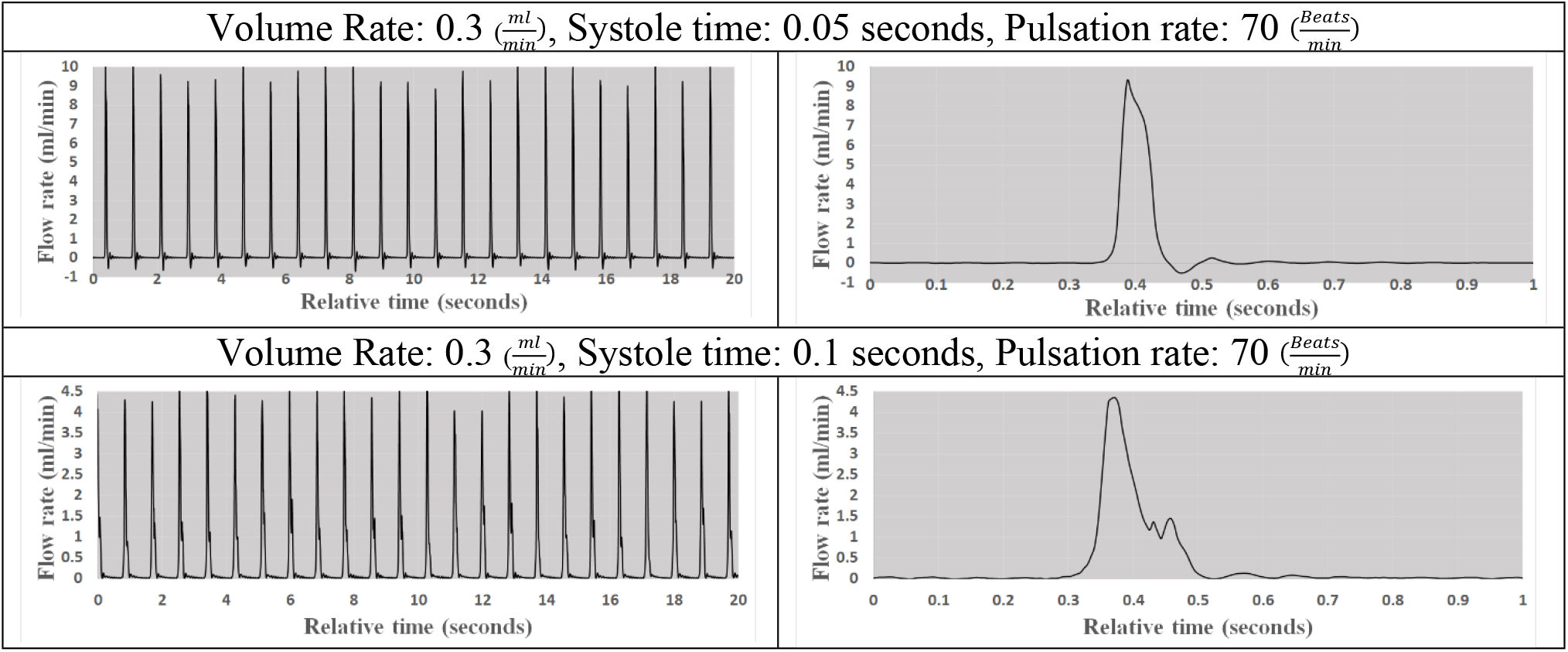
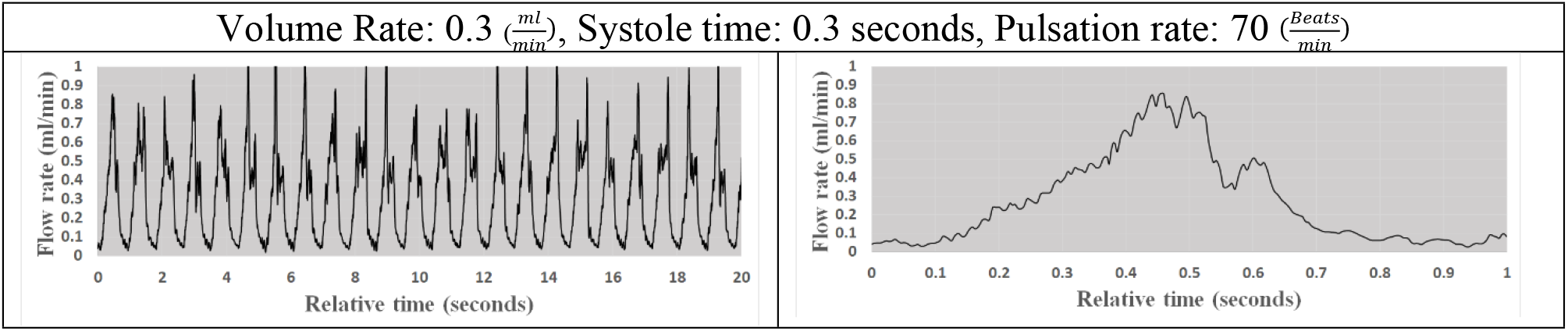
Visual representation of flow pattern at various systole times while heart rate and volume rate remained constant at 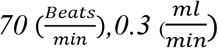,

In the first set of flow patterns, shown in Figure 4, the beat rate and systole time remain constant at 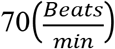 and 0.1 seconds while the output volume rate varies between 0.35, 0.1, 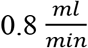. These results indicate that the pumps are capable of variable output regardless of beat rate or systole time.

In Figure 5. systole time and output volume rates remained constant at 0.1 seconds and 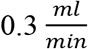 while the beat rate changed from 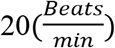 to 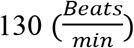 and 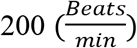. The number of individual peaks in each 20-second segment in Figures correctly corresponds to the specified beat rate.

Finally, in Figure 6, the beat rate is maintained at 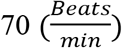 and the output volume rate remain constant at 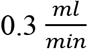 while systole time varies from 0.05 to 0.1 and 0.3 seconds. As illustrated in Figures 2 and Figure 4, the pump can maintain a specified beat rate independent of systole time and volume rate.

## Discussion

Using the data acquisition system described earlier, sample flow rates were collected over 20-second intervals to demonstrate the pumps’ capabilities in recreating pulsatile flow at physiologically accurate CSF production rates. Variation testing across the three input parameters used to define CSF production rate, those being volume rate, heart rate, and systole time, was also performed to demonstrate the effect of these variables on the subsequent flow pattern. Because of the dampening effects of the tubing and valve system, amplitudes of these flow patterns are reduced across the range of the 20-second interval. Furthermore, zooming in on an individual pulse during the 20-second interval, it is evident that the pump induces an initial rise in flow rate typical to flow patterns recorded during a normal cardiac cycle, followed by a constant reduction in flow rate leading up to the next pulse in the profile.

For this study, a consistent silicone tubing setup described earlier was used throughout the testing. The manipulation of this tubing set up in the future and in turn, the dampening factor applied across the individual channels of the pump as well as the pump will not only manipulate the amplitude of each beat within the profile as described by Equation 1 but can also affect the minor spike found on the downward stroke of each beat cycle.

The emphasis on the manipulation of the controllable variables of the pump is essential to the success of the pump across many disciplines where accurate tabletop recreations of flow rates and patterns are currently a limiting factor. Specifically for hydrocephalus, the instances presented here recreate the physiologic flow rates and patterns for CSF production which is particularly important as control groups to be compared to the clinical state in future studies.

It has been reported that the fluid dynamics of CSF production and absorption are influenced by a broad spectrum of factors such as the individuals’ unique anatomy and physical characteristics as well as real-time physiologic conditions such as blood pressure and heart rate [22–25]. Furthermore, the input from the central nervous system adds another layer of complexity and variability in CSF dynamics. Therefore, optimal operation over a reasonable range relative to the physiologic conditions was one of the main objectives in the design and development of this device. One of the main advantages of this device over previous models is flexibility in the domain of operation (Table 3). A mean CSF production rate of 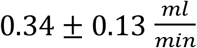 was reported by Edsbagge et. al [26]. Profiles at average physiologic heart rate and systole time were used to test the capabilities of the pump recreating the mean reported CSF production rate along with the upper and lower limits of the mean. The domain shown within Table 3 for physiologic CSF production rate has shown to be accurate across its range.

**Table 3.** Volumetric analysis of pumps running within the mean CSF production rate reported by Edsbagge et. al [26]

| Input | Output |  |  |  |  |
| --- | --- | --- | --- | --- | --- |
| Volume Rate ($\frac{ml}{min}$) | Mean of 15 channels ($\frac{ml}{min}$) | Lowest Value ($\frac{ml}{min}$) | Highest Value ($\frac{ml}{min}$) | Mean Absolute Error ($\frac{ml}{min}$) | Mean Relative Error |
| **0.21** | 0.209015533 | 0.202983 | 0.211548 | 0.000984 | 0.471002% |
| **0.34** | 0.345996 | 0.33252 | 0.350533 | -0.006 | 1.733025% |
| **0.47** | 0.462307 | 0.45535 | 0.467528 | 0.007693 | 1.664075% |
| **Overall mean:** |  |  |  | 0.000892 | 1.289367% |

The validation of this tabletop model specifically for the use in isolated anatomical subdivisions of human production and absorption system of CSF is limited because of the current difficulties in measuring ICP and CSF flow. However, because of the modular tubing setup of the system, additions to the tube setup in series allow for the easy testing of future components essential to the modern treatment of hydrocephalus and other domains of research such as catheters and valves. The scale of the pump and tubes along with the incorporation of other components such as bioreactor chambers make it a much simpler system to work with within a sterile environment where an incubator and live cells are involved. Furthermore, it has been reported that compliance may have a major physiologic function in CSF flow characteristics and distribution [19,27]. Therefore, future studies will also investigate the addition of compliance components to the pump. This study demonstrated that this setup could maintain a CSF production rate, beat rate, and systole time profile for an extended period over 15 channels. However, these variables all change throughout the day not only in hydrocephalic patients but in healthy individuals as well [28–30]. Future iterations of the pump and the controlling program can integrate a system where patient data can be recreated across a full day. The implications of a system like this could alter the perspectives of researchers within this field and other fields where this pump is applicable, where an all-encompassing system may be the new standard in terms of accurate in-vitro testing.

## Conclusion

In conclusion, the validation of this reciprocating positive displacement pump system allows for the future validation of novel designs to biomaterials and devices used in the treatment of hydrocephalus. Specifically, the use of this system includes major components such as a hydrocephalus catheter and chamber capable of accurately reflecting changes in ventricular expansion, as well as a hydrocephalus valve used to regulate the flow of CSF.

## Declarations

### Authors’ Contributions

JG: Conceptualization, Methodology, Formal analysis, Investigation, Data curation, Project administration, Writing—original draft, Visualization, and Writing—review & editing.

